# Comparing Orientation-Dependent Transverse Relaxation at 3 T and 7 T: Deciphering Anisotropic Relaxation Mechanisms in White Matter

**DOI:** 10.1101/2025.05.05.652295

**Authors:** Yuxi Pang, Rajikha Raja, Wilburn E. Reddick

## Abstract

Purpose: This work aims to elucidate the mechanisms underlying orientation-dependent transverse relaxation in human brain white matter (WM), which have long been ambiguous. Methods: We analyzed publicly available 3T and 7T DTI datasets (b-values = 1000 and 2000 s/mm2) from 25 young adults participating in the Human Connectome Project. Orientation-dependent transverse relaxation R2 profiles from whole brain WM were generated from T2-weighted images (b-value = 0) and characterized using a previously developed cone model based on the generalized magic angle effect (MAE). Derived anisotropic R2 or R2a values were compared between different magnetic fields. Similar comparisons were also conducted for the derived R2a values from whole brain WM and two fiber tracts, based on gradient-echo signals reported in the literature. Classical relaxation theories predict that the ratio of (R2a (7T))/R2a (3T)) will be unity or (7/3)^2 (i.e., approximately 5.4) if the measured orientation-dependent R2 arises exclusively from MAE or from the previously proposed susceptibility effect, respectively. Results: Fitted model parameters were comparable for DTI datasets with b-values of 1000 and 2000 (s/mm2). The fitted R2a increased, on average, from 3.6±1.1 (1/s) at 3T to 5.4±1.5 (1/s) at 7T for DTI datasets with a b-value=1000 s/mm2. The measured ratio of (R2a (7T))/(R2a (3T)) was thus approximately 1.5. However, based on gradient-echo signals, this ratio essentially became unity within the measurement precision. Conclusion: This study suggests that MAE is the primary mechanism for the observed orientation-dependent transverse relaxation at 3T in human brain WM, offering a different perspective from previous literature.

## 1 INTRODUCTION

The distinct contrasts observed in magnetic resonance (MR) imaging of biological tissues are strongly influenced by the composition of biomolecular constituents and their hierarchical organization [1, 2]. Multiple relaxation mechanisms—including (residual) dipolar interactions, chemical exchange, and (anisotropic) susceptibility effects—contribute to these image contrasts. For example, both susceptibility-based pathways and the magic angle effect (MAE) have recently been considered [3] for characterizing orientation-dependent transverse relaxation in brain white matter (WM). However, MAE has not been adequately addressed, largely due to an oversight of the detailed organization of bound water adjacent to or within biomacromolecules. Historically, the concept of “structured” water in tissues has been met with skepticism [4]; nevertheless, its recent application via MAE to orientation-dependent transverse relaxometry in neural tissues [5, 6] has highlighted the role of microstructural water organization, challenging previously prevalent susceptibility-based models [3, 7, 8].

Approximately two decades ago, the increasing availability of whole-body MR scanners operating at ultra-high magnetic *B*_0_ fields led to the discovery of unprecedented varying 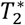 contrast in gradient-echo (GRE) images of human brain WM in vivo at 7 T [9, 10]. This characteristic 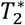 contrast (or its reciprocal, 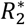 contrast) was subsequently found to depend on the orientation of axon fiber tracts in an animal study [11], where the anesthetized brain was positioned at different angles relative to the *B*_0_ field. Shortly thereafter, a clearer picture emerged from a study conducted on healthy human brains at 3 T [12], where the 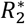 values in fiber tracts were found to be higher along either the left-right or anterior-posterior directions compared to the inferior-superior direction, which aligns with the *B*_0_ field. Moreover, the observed 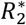 values appeared to be proportionally correlated with diffusion fractional anisotropy (FA) derived from diffusion tensor imaging (DTI).

The complete profiles of 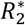 orientation dependence in human brain WM in vivo at 3 T were eventually established by leveraging orientation information from DTI, specifically using the principal diffusivity (*λ*_1_) direction within each imaging voxel [13, 14]. More importantly, an orientation-dependent function of sin^2^ *θ*(or equivalently sin 2*θ* [15]) was proposed, where *θ* is the angle between the *λ*_1_ direction and the *B*_0_ field. In a subsequent study involving two coronal formalin-fixed brain slabs at 7 T [16], an enhanced orientation-dependent function was developed, incorporating an additional sin 4*θ* term. This refined function was derived from 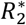 measurements of brain specimens rotated at 18 different angles around the brain slab’s norm direction, spanning from 0° to 170° relative to the *B*_0_ field. In this ex vivo study, the investigators also introduced a phase shift (*φ*_0_) into the proposed 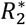 orientation-dependent function. However, the biophysical basis of *φ*_0_ was not adequately elucidated. As a result, this phase shift was often overlooked in subsequent applications of the function to previously reported studies, where orientation information was derived from *λ*_1_ in DTI [17-19].

To better understand the origins of the observed orientation-dependent transverse relaxation in WM, a follow-up investigation was conducted at 7 T using rotating brain specimens, following a workflow similar to that described above [3]. In this comparative study, unlike the pronounced 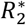 orientational anisotropy, the measured *R*_2_ values exhibited minimal, if any, discernible orientation dependence. This observation led investigators to conclude that magnetic susceptibility effects were the dominant mechanism underlying anisotropic relaxation. However, the temperature-dependent anisotropies of 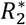 observed in the brain samples could not be adequately explained by the proposed relaxation mechanism, because the diamagnetic susceptibility of myelin is essentially independent of temperature and the static dephasing theory invoked therein also neglects spin motion. These conflicting findings clearly illustrate that the true origin of orientation-dependent 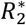 in WM remains elusive.

Meanwhile, researchers developed a hollow cylinder fiber model (HCFM) for WM based on magnetic susceptibility theory [7, 20]. In this model, the annular region of the cylinder is filled with molecular constituents exhibiting anisotropic magnetic susceptibilities, allowing prediction of 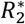 orientation dependence using an analytical function that includes a sin^4^ *θ* or sin 4*θ* term [7]. However, this analytical function alone was insufficient to accurately characterize the shape of orientation-dependent 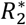 profiles measured from rotating pig brain samples, unless an additional heuristic sin^2^ *θ* term was included. Interestingly, a similar orientation-dependent *R*_2_ function, which incorporates the term sin^4^ *θ*, with and without the term sin^2^ *θ*, was also proposed for human brain WM in vivo at 3 T [8]. The underlying biophysical principle of the proposed model for *R*_2_ orientational anisotropy was the same susceptibility theory for 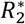 anisotropy but applied at a mesoscopic scale. Specifically, diffusion-mediated decoherence, arising from local susceptibility variations across axon fiber bundles, enhances the transverse relaxation measured at the macroscopic voxel level.

Another potential anisotropic relaxation mechanism is the well-known MAE [21], originated from rotationally restricted water molecules in highly ordered biological tissues [22], but this effect has not been adequately considered in existing orientation-dependent 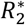 or *R*_2_ relaxation models [16, 23]. This oversight is partially due to the markedly different profiles of measured 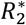 or *R*_2_ orientation dependence in WM, where enhanced relaxation rates are observed when axon fibers are oriented perpendicular to the *B*_0_ field, compared to parallel [12]. In contrast, conventional MAE profiles, such as those observed in tendons and deep-zone cartilage, exhibit the opposite pattern of transverse relaxation rate enhancement between these two orientations [21]. Recently, this seemingly contradictory 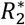 or *R*_2_ anisotropy in WM has been reconciled by employing a generalized MAE function that incorporates a phase shift [5, 6]. Remarkably, the proposed MAE-based function can be reformulated in terms of previously developed orientation-dependent 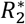 and *R*_2_ models, thereby suggesting a potentially ambiguous mechanism underlying anisotropic transverse relaxation in WM, as reported in prior literature [3, 15].

An orientation-dependent transverse relaxation rate provides essential specificity in neuro-microstructural imaging, as it directly reflects the extent of (de)myelination in WM [3, 5, 20, 24]. To accurately capture the underlying biophysical processes, it is essential to disentangle the previously proposed relaxation mechanisms—namely, the susceptibility effects and the MAE—in relation to 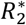 and *R*_2_ orientational anisotropies. These two competing anisotropic relaxation pathways exhibit distinct *B*_0_ field dependences: susceptibility effects scale with 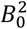, whereas the MAE is independent of *B*_0_. Consequently, these mechanisms can, in principle, be distinguished by comparing orientation-dependent *R*_2_ across different *B*_0_ field strengths. This separation is critical for advancing our understanding of the unique anisotropic relaxation signatures associated with the highly organized microstructures in WM. Given the current ambiguity surrounding the mechanism of orientation-dependent transverse relaxation, this work aims to clarify the long-standing question regarding the origins of anisotropic transverse relaxation in human brain WM by examining orientation-dependent *R*_2_ values derived from both spin-echo and gradient-echo measurements at 3 T and 7 T, as previously published in the literature. Compared to our prior investigations [5, 6], this study represents a significant advance by identifying the dominant anisotropic relaxation pathway at clinically widely available *B*_0_ field strengths.

## 2 THEORY

This section provides a brief overview of previously developed biophysical models describing the orientation dependence of 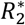 and *R*_2_ in brain WM. The literature indicates that the orientation dependence of 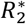—which equals 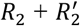—originates primarily from *R*_2_ rather than 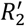 [7, 25]. It is generally accepted that 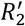 arises mainly from non-local effects caused by large length-scale *B*_0_ inhomogeneities and macroscopic variations in magnetic susceptibility [7]. Because this non-local contribution is time-independent, it can be removed during post-processing when deriving the orientation dependence of 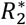.

Minor anisotropic 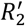 contributions may arise from WM boundaries; however, the data-binning workflow used in this study would likely reduce their potential impact. Nevertheless, this issue should be carefully considered in ex vivo studies of brain WM samples. Accordingly, in the present study, 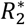 and *R*_2_ are treated interchangeably with respect to orientation dependence, even though they are derived from GRE and spin-echo (SE) datasets, respectively. This approach is consistent with previous reports [6, 25].

### 2.1 Susceptibility-based models for 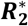 and *R*_2_ orientation dependences

According to the previously proposed HCFM model [7, 20], an orientation-dependent function for 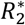 is given in Equation 1, which includes an analytically derived term sin^4^ *θ* and a heuristic term sin^2^ *θ*. The latter term is essential for accounting for myelin water contributions to both simulated and observed 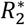 values in WM [7]. A seemingly different, yet mathematically equivalent [15], orientation-dependent function for 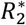 has also been developed in the literature [3], as shown in Equation 2:

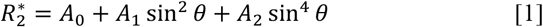

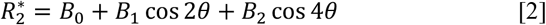

Although mathematically equivalent, these two functions arise from fundamentally different biophysical assumptions. The sin^2^ *θ* and sin^4^ *θ* terms in Equation 1 originate from myelin water in the static dephasing regime and from intra- and extra-axonal water in the diffusion or motional narrowing regime, respectively [7]. In contrast, Equation 2 was derived under the assumption that the measured signals are entirely in the static dephasing regime [26], where translational diffusion is negligible. The cos 2*θ* and cos 4*θ* terms are attributed to the highly ordered myelin microstructure and its associated susceptibility anisotropy [3].

Interestingly, Equation 1 has also been proposed to characterize the orientation dependence of *R*_2_ [8], where the sin^4^ *θ* term arises from diffusion-mediated decoherence due to mesoscopic magnetic susceptibility differences, and the sin^2^ *θ* term results from interactions between susceptibility differences and applied field gradients. While the latter contribution is expected to be negligible in standard SE datasets, it appears necessary to include the sin^2^ *θ* term to better fit the orientation dependence profile of *R*_2_ in WM [15, 19].

### 2.2 A generalized orientation-dependent model derived from MAE

Given the conflicting biophysical foundations of previously developed models, a generalized orientation-dependent function has been proposed for WM that relies solely on the well-established MAE [5, 6]. As schematically illustrated in Figure 1A (adapted from the literature [27]), a typical myelinated axon is wrapped in multiple layers of bilayer membranes. Rotationally restricted water can reside both within an axonal space [17, 28], denoted by a filled violet circle, and in the compartments containing myelin water [29], indicated by the label H2O (Figure 1B). In the latter case, the *intramolecular* proton-proton (H-H) residual dipolar interaction (RDI) vectors from rotationally restricted water are assumed to lie within the plane perpendicular to the axon axis (Figures 1A-1C) [5]. In contrast, the RDI vectors within the axonal space are assumed to align along the axon’s longitudinal axis [17]. These two extreme cases correspond to the RDI vector distributions in ideal collagen fiber organizations in the superficial (tangential) and deep (radial) zones [30, 31].

**FIGURE 1.**
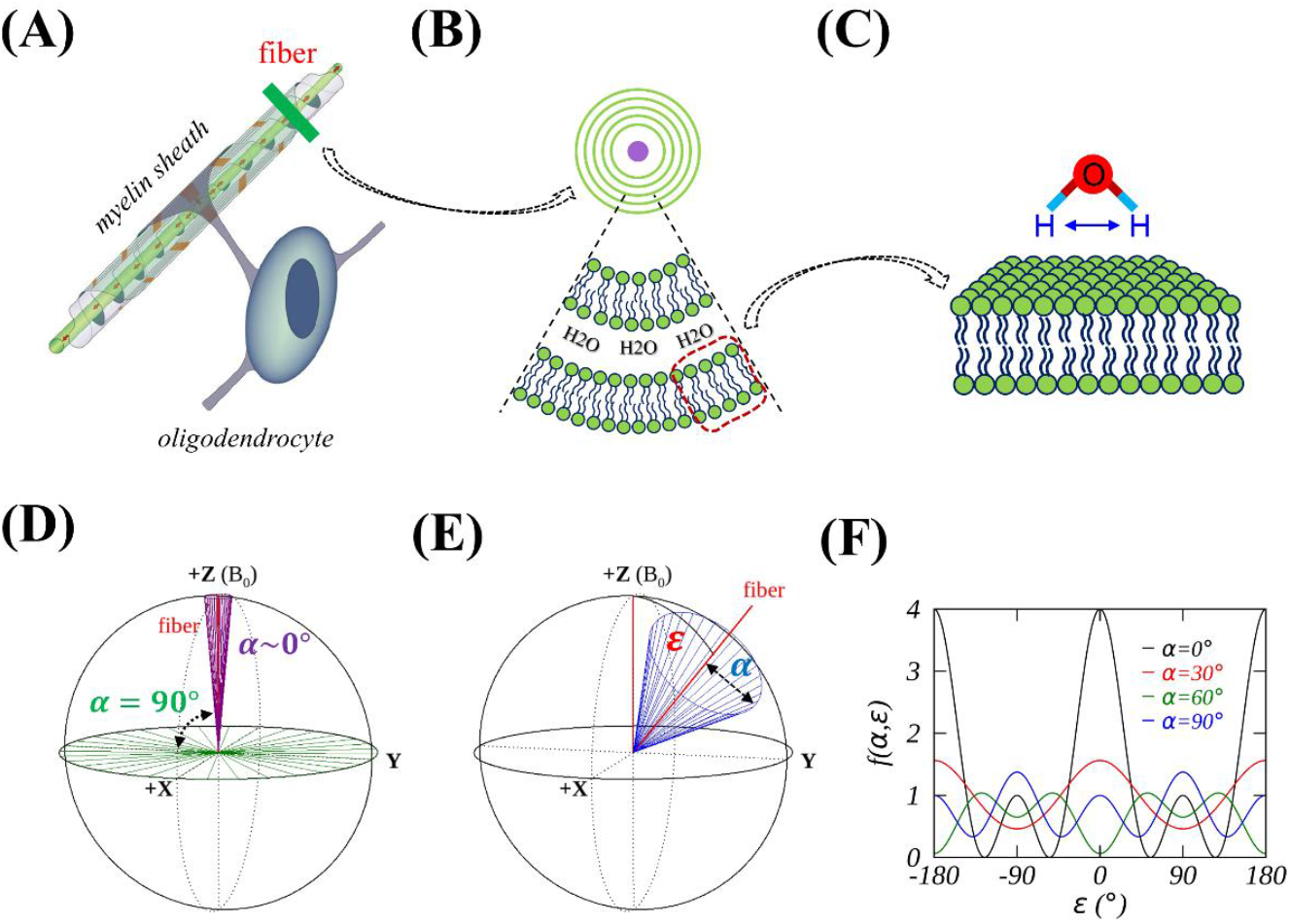
Schematics illustrate the hierarchical structures of rotationally restricted water in an axon fiber and a cone model for orientation-dependent transverse relaxation. (A) Myelinated axon fiber, adapted from Reference 25. (B) Cross-sectional view perpendicular to the fiber, showing restricted water located within the axon (filled violet circle) and in the extra-axonal space (between membrane bilayers). (C) Restricted water molecule near the membrane surface. (D) Restricted water within the axon (α=0°, violet) and outside the axon (α=90°, green). (E) Cone model with an opening angle α, orientated at an angle ε relative to the static magnetic field B_0_. (F) Theoretical orientation dependence of transverse relaxation. Note: when referring to the orientation of restricted water, this denotes the direction of the residual dipolar interaction between two protons within the water molecule, as schematically illustrated in panel (C).

A generalized function was derived by evaluating the effective RDI vectors within an axially symmetric system, modeled as a cone (Figure 1E) with an opening angle *α*. This angle represents the relative contributions from water inside (*α* = 0°) and outside (*α* = 90°) the axon (Figure 1D) [5]. In this model, the effective RDI vectors, characterized by the orientation-dependent term (3cos^2^*θ* − 1)^2^, are distributed along the cone wall. If the cone’s axis forms an angle *ε* with respect to the *B*_0_ field, the orientation dependence of *R*_2_ from this ensemble of RDI vectors can be concisely expressed as [5, 6]:

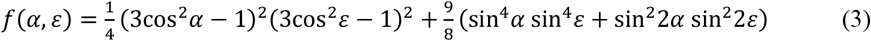

It has previously been shown that Equations 1-3 are mathematically equivalent, differing only in their scaling factors [6, 15]. When *α* = 0°, this generalized function reduces to the conventional MAE expression: *f*(0°, *ε*) = (3cos^2^*ε* − 1)^2^. Conversely, when *α* = 90°, this function becomes: *f*(90°, *ε*) = (1/4)(3cos^2^*ε* − 1)^2^ + (9/8)(sin^4^*ε*). As shown in Figure 1F, the orientation-dependent profile of *f*(0°, *ε*) (black curve) differs markedly from that of *f*(90°, *ε*) (blue curve). Confusion may arise if the observed orientation-dependent measures in WM are interpreted using the conventional MAE expression.

The transverse relaxation rate *R*_2_ measured in WM can generally be decomposed into two components: an orientation-independent term 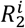, and an orientation-dependent term 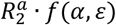, expressed as:

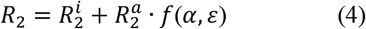

In this formulation, 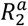 represents the magnitude of the orientation dependence, which reaches its maximum value of 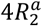 when both *α* and *ε* are zero [6].

When considering an effective transverse relaxation rate 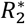, defined as 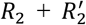, the contribution from 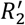 is incorporated into the orientation-independent term 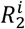. It is important to note that 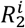 is typically assumed to be constant when analyzing orientation-dependent phenomena in biological tissues.

### 2.3 Orientation dependence of *R*_2_ derived from a single *T*_2_-weighted image

As shown in the literature [5, 6], 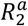 can be extracted from T2W images when voxel-wise fiber orientations (*ε*) are available. The signal of a typical voxel (*S*_*b*=0_) in a T2W image can be expressed as

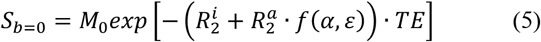

Here, *M*_0_ is the maximum signal amplitude as *TE* approaches zero. In logarithmic form, Equation 5 can be rewritten as

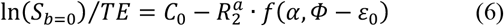

where *C*_0_ is an unknown constant defined as 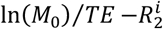, *ε*_0_ is a phase shift [6], and *Φ* denotes the principal diffusivity direction relative to the *B*_0_ field (see Section 3.1).

As previously reported, *M*_0_ varies minimally across WM throughout the brain [32] and is therefore treated as constant. It is worth noting that the definition of *M*_0_ may be susceptible to some *T*_1_ effects, which have also been reported to be orientation-dependent [33]. However, these potential effects are negligible, as the degree of *R*_1_ (=1/*T*_1_) anisotropy is at least two orders of magnitude smaller than that of *R*_2_ anisotropy.

### 2.4 *B*_0_-dependence of anisotropic transverse relaxation rates

As shown above, Equations 1-3 are mathematically equivalent when scaling factors are not considered, yet they are derived from distinct biophysical foundations. Therefore, it is not feasible to disentangle two different relaxation mechanisms in WM merely by rearranging orientation-dependent terms in previously proposed model functions [3, 15]. If magnetic susceptibility were the sole mechanism driving orientation-dependent relaxation, then 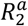 would be expected to scale proportionally with 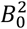 [7, 8]. Conversely, if MAE were responsible for anisotropic transverse relaxation, then 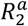 would be entirely independent of *B*_0_ [22, 34].

Accordingly, these two relaxation mechanisms can be disentangled by comparing 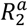 values measured at two different magnetic field strengths, as demonstrated in the literature [35]. For example, the ratio between 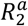 measured at 7 T and 3 T, denoted by *η*, is theoretically predicted to be *η* = (7/3)^2^ = 5.44, assuming the increase in 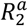 is solely due to susceptibility effects. When considering the actual *B*_0_ field strengths used in data acquisitions [36], i.e., 2.895 T and 6.980 T, *η* increases slightly to 5.81. This more realistic value is used in the following data analysis, although the field strengths will continue to be referred to as 3 T and 7 T for convenience.

In a different scenario, susceptibility may partially contribute to the observed 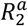 in WM. In this case, the percentage contribution of susceptibility to the measured 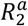 at 3 T can be quantified as described in the literature [35]: 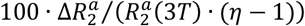, where 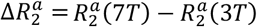. Similarly, the partial contribution to the measured 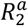 at 7 T can be evaluated as: 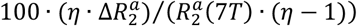. Nonetheless, any conclusion regarding the relative importance of specific anisotropic transverse relaxation mechanisms should remain consistent, regardless of whether 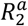 is measured using GRE or SE signals.

## 3 METHODS

### 3.1 Publicly available DTI datasets at 3 T and 7 T from HCP

After signing the data usage agreement, we randomly selected 25 participants from a cohort of 184 young adults in the Human Connectome Project (HCP) [https://www.humanconnectome.org/study/hcp-young-adult]. Each subject had DTI datasets acquired at both 3 T (TE = 89 ms) and 7 T (TE = 71 ms). As shown in Tables S1 and S2, each subject is identified by a subject number (1^st^ column) used in the present study, along with their HCP-ID (2^nd^ column), which specifies the original HCP identification number, gender, and age. For example, the 15th subject’s HCP-ID is 385046, the gender is male (M), and the age is 28 years.

These publicly available DTI datasets had already been preprocessed to correct image artifacts caused by signal drift, susceptibility effects, eddy currents, gradient nonlinearity-induced distortions, and involuntary head motion. Data subsets with *b-*values of 0 and 1000 s/mm^2^, as well as *b-*values of 0 and 2000 s/mm^2^, were separately fitted using FSL’s DTIFIT software [37]. In other words, each participant contributed two datasets at a specific field strength. The fitted parameters included three eigenvalues (*λ*_1_, *λ*_2_, *λ*_3_), three eigenvectors (ê_1_, ê_2_, ê_3_), fractional anisotropy (FA), mode of anisotropy (MO) [38], and *T*_2_-weighted (T2W) images. The direction *Φ* of the principal diffusivity *λ*_1_, relative to the *B*_0_ field, was represented by the primary eigenvector ê_1_ using the relationship [13, 19]: 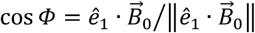.

WM voxels were segmented from high-resolution anatomical *T*_1_-weighted volumetric images in the HCP datasets. These voxels were further refined using thresholds of 0.5 < FA < 0.9 and 0.5 < MO < 1.0, ensuring that each WM voxel was well-characterized by a prolate diffusion tensor [6]. Note that *M*_0_ and MO refer to distinct entities. Figure 2 shows the final WM masks superimposed on T2W images from DTI acquired at 3 T and 7 T. After pooling and binning data from the selected WM voxels across the brain, orientation-dependent profiles of 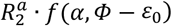, defined as *C*_0_ − In(*S*_*b*=0_)/*TE*, were established as demonstrated in prior studies [5, 6]. The four model parameters 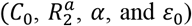 were extracted using a nonlinear least-squares curve fitting method [39]. In total, fifty orientation-dependent *R*_2_ profiles were generated, with each participant contributing two datasets corresponding to distinct non-zero *b-*values at a specific field strength; this sample size allows reliable conclusions to be drawn.

**FIGURE 2.**
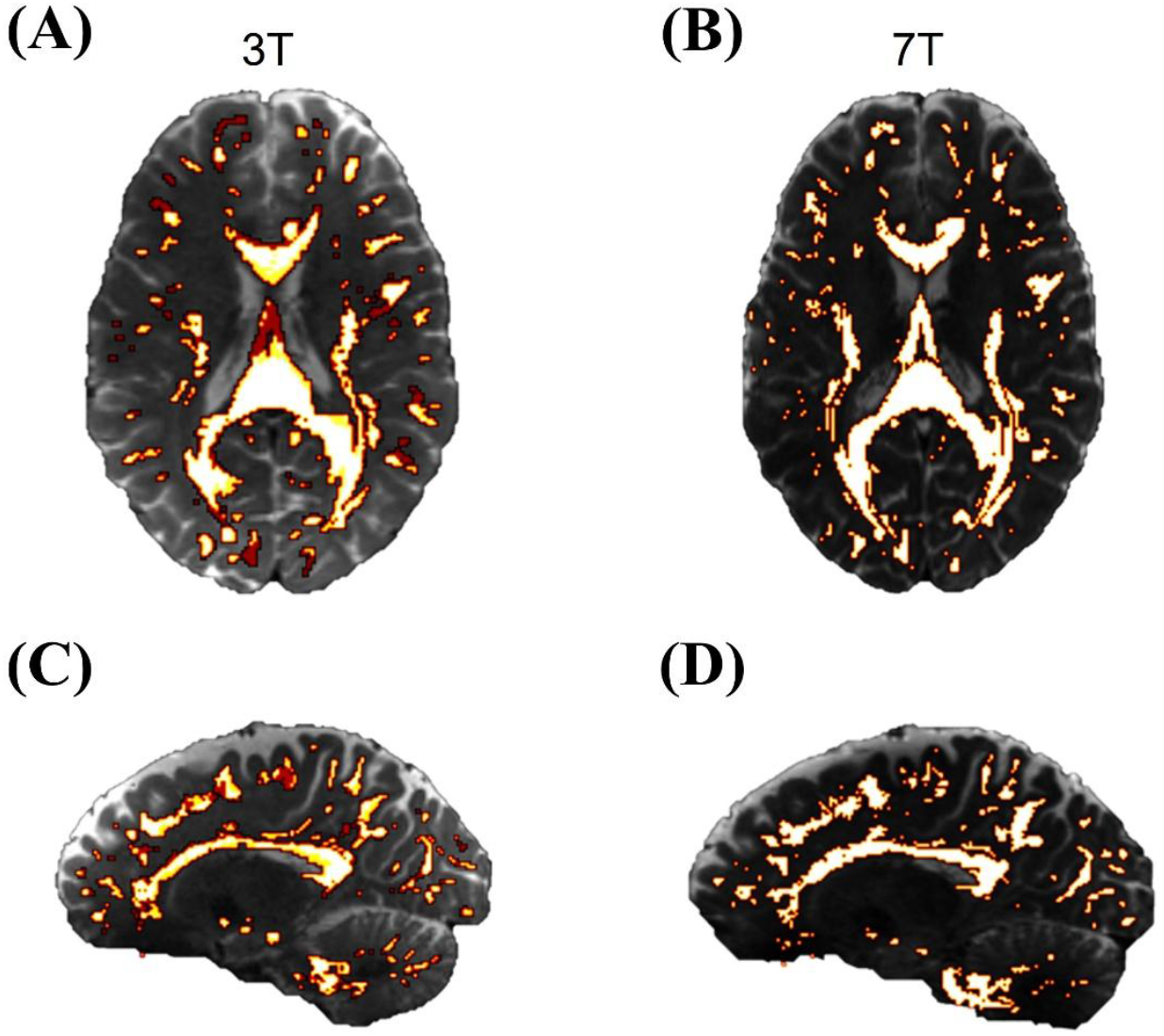
White matter (WM) masks overlaid on T_2_-weighted (b-value = 0) images in axial (A, B) and sagittal (C, D) views from 3 T (A, C) and 7 T (B, D) DTI datasets. Although the WM masks are displayed using a continuous colormap for visualization purposes, the underlying data are binary (0 or 1) and were used as such in all analyses.

### 3.2 Previously published orientation-dependent 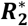 profiles at 3 T and 7 T

A free online tool (www.graphreader.com) was used to extract previously published orientation-dependent 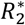 profiles of WM at 3 T and 7 T. These profiles were measured either across whole-brain WM or within specific fiber tracts in healthy subjects. The reported 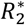 datasets were acquired in accordance with ethical guidelines, as stated in the original publications.

The first 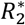 profile of whole-brain WM at 3 T [40], free from the confounding flip angle effect (FAE), was derived by linear regression from profiles measured at the flip angles of 6°, 17°, 35° and 60°, as described in the literature [41]. This orientation-dependent 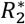 profile at 3 T, independent of FAE, was averaged across six subjects (age 22-29; median age 25). The second 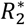 profile of whole-brain WM at 7 T, also free from FAE, was extracted directly from Figure 7 (Session2, green) of the original paper [41]. This profile was measured in three healthy subjects (mean age 38.7 ± 4.2 years).

The third and fourth 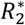 profiles were derived from the corpus callosum (CC) at 3 T and 7 T, respectively, while the fifth and sixth profiles were obtained from the cingulum (CG) at 3 T and 7 T [42]. These 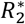 profiles were reported in a conference abstract that did not include specific demographic information. However, a related publication based on partial datasets indicated that five healthy participants were involved, with an average age of 34 ± 6 years [43].

It is worth noting that the orientation information underlying the 7 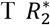 profiles was derived from 3 T DTI data [41, 42]. Furthermore, the orientation used to describe all 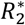 profiles, except for the second, was defined as the angle between the *B*_0_ field and the direction of the first peak of the orientation distribution function (ODF), rather than the direction (*Φ*) of the principal diffusivity, which is more commonly used. The ODF was previously computed using the constrained spherical deconvolution method applied to multi-shell diffusion datasets [44]. For consistency, these ODF-based angles are still denoted as *Φ* in this work.

### 3.3 Data analysis and error estimations

Orientation-dependent 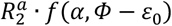 profiles based on single SE signals were generated using custom Python scripts. Each profile consisted of 18 data points, covering orientations (*Φ*) from 2.5° to 87.5°. A publicly available nonlinear least-squares fitting routine (http://purl.com/net/mpfit) [39], written in Interactive Data Language (IDL 9.1, Harris Geospatial Solutions, Inc., Broomfield, CO, USA), was used to characterize these orientation-dependent profiles based on Equation 6. For profiles derived from GRE signals, the fitting was performed using Equation 4, where the theoretical fiber orientation *ε* was replaced with *Φ* − *ε*_0_.

The fitted model parameters were constrained to the following ranges:

1. SE-based profiles: *C*_0_ = [0, 200] s^-1^ and 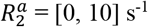
2. GRE-based profiles: 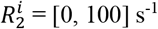 and 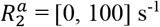
3. All profiles: *α* = [0°, 90°] and *ε*_0_= [0°, 45°]

Note that 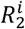 could not be determined from T2W images acquired with only one TE, as is typically the case in standard DTI datasets. Because the nonlinear least-squares fittings were unweighted, the reported uncertainties, formal 1σ errors in each fitted parameter derived from the covariance matrix, were rescaled such that the reduced χ^2^ was approximately unity. More information can be found in the commentary section of the fitting routine (i.e., mpfit.pro).

Unless otherwise stated, error propagation was performed according to the standard procedures as described in [45]. Standard deviations of the presented data are indicated by error bars or by the half-widths of shaded ribbons. Image data were visualized using the MRtrix3 viewer (MRView; Figure 2) [46].

## 4 RESULTS

### 4.1 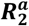 derived from SE signals at 3 T and 7 T

Figure 3 compares representative orientation (Φ) dependence profiles of 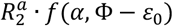 between 3 T (blue) and 7 T (red) from the 15^th^ subject (HCP_ID: 385046_M28). These profiles were derived from non-zero *b*-value of 1000 s/mm^2^ (Figure 3A) and 2000 s/mm^2^ (Figure 3B). For the 1000 s/mm^2^ data, the fitted 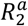 increased from 4.16 ± 0.39 s^-1^ at 3 T to 6.27 ± 0.58 s^-1^ at 7 T, while other fit parameters, such as *α* and *ε*_0_, remained comparable. Comparing Figure 3B with Figure 3A, the fitted 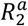 appears relatively larger at the higher *b*-value. This increase is attributable to the fixed selection thresholds for FA and MO used in this study, as well as the inverse relationship between *b*-value and FA, as discussed in our prior work [6]. The ratio *η* of 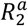 values between 7 T and 3 T was 1.51 ± 0.20. For the 2000 s/mm^2^ data, this ratio (*η* = 1.60 ± 0.25) was nearly identical, within measurement uncertainties. Notably, these measured *η* values are significantly lower than the theoretical ratio (*η* = 5.81) expected if susceptibility effects alone accounted for the observed orientation-dependent transverse relaxation in WM.

**FIGURE 3.**
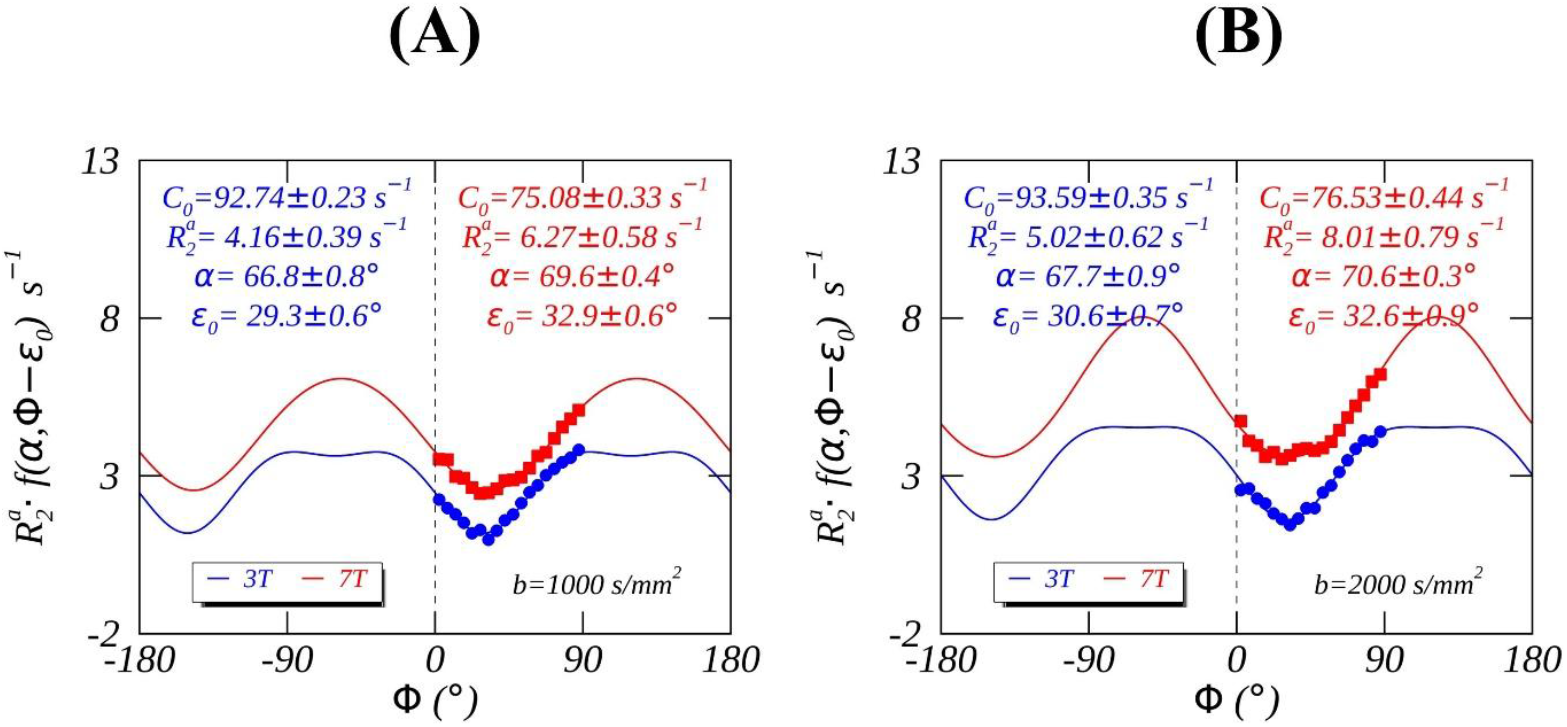
Measured (colored symbols) and fitted (colored lines) orientation-dependent transverse relaxation rates 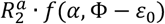 from Subject 15 at 3 T (blue) and 7 T (red), based on DTI using non-zero b-values of 1000 s/mm^2^ (A) and 2000 s/mm^2^ (B).

Across all 25 subjects included in this study, the fitted 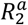 values at 3 T are plotted against those measured at 7 T in Figure 4, for datasets with *b*-values of 1000 s/mm^2^ (Figure 4A) and 2000 s/mm^2^ (Figure 4C). In these plots, colored dashed lines represent two theoretically predicted *η* values corresponding to exclusive anisotropic relaxation mechanisms: susceptibility effects (*η* = 5.81, red) and MAE (*η* = 1.0, blue). Subject-specific *η* values are shown in Figure 4B and Figure 4D, with mean values of 1.39 ± 0.50 and 1.47 ± 0.40, respectively, indicated by dashed horizontal lines. Detailed fitted parameters are provided in Table S1 (*b*-value=1000 s/mm^2^) and Table S2 (*b*-value=2000 s/mm^2^).

**FIGURE 4.**
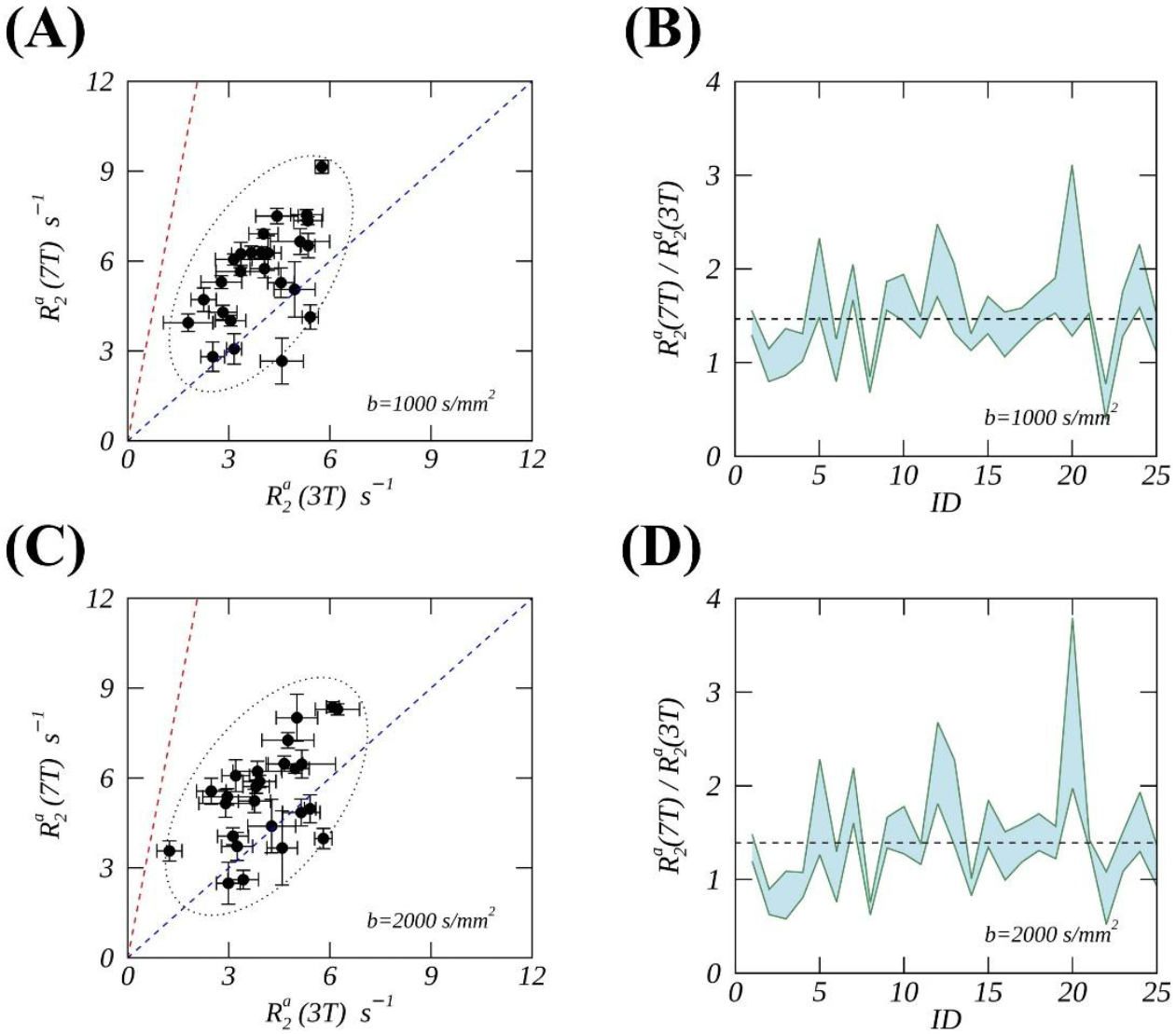
Scatterplots (A, C) and ratio plots (B, D) of derived 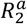 from 7 T and 3 T DTI datasets across 25 subjects. DTI datasets were obtained using non-zero b-values of 1000 s/mm^2^ (A, B) and 2000 s/mm^2^ (C, D). In the scatterplots, colored dashed lines represent the theoretical ratios of 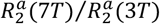 assuming anisotropic transverse relaxation arises from susceptibility effects (red) or the magic angle effect (blue). In the ratio plots, horizontal dashed lines indicate the average ratios measured across the 25 subjects. Note that standard deviations are indicated by error bars (A, C) and by the half-widths of the shaded ribbons (B, D). The ellipses in panels A and C represent the 95% confidence level.

From 3 T to 7 T, as shown in Table S1, the fitted 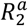 values increased on average from 4.0 ± 1.1 s^-1^ to 5.6 ± 1.6 s^-1^, corresponding to 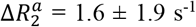. The percentage contribution of susceptibility effects to the measured 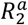 at 3 T was 8.3 ± 10.2%, compared with 34.5 ± 42.2% at 7 T. Comparable results were obtained from the fits in Table S2; specifically, the susceptibility-based contributions at 3 T and 7 T were 6.6 ± 10.3% and 29.1 ± 45.6%, respectively,

These results suggest that MAE predominantly contributes to the measured 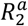 at 3 T, consistent with recent findings in the literature [25, 47]. As shown in Table S1, the average fitted *α* increased from 66.5 ± 2.3° to 69.6 ± 1.8°, while *ε*_0_ decreased from 26.2 ± 3.8° to 11.5 ± 11.0°. These fitted model parameters and their relative changes from 3 T to 7 T are consistent with those reported in Table S2.

### 4.2 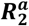 derived from GRE signals at 3 T and 7 T

When comparing the fitted parameters from the orientation-dependent 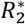 profiles of whole-brain WM at 3 T (Figure 5A) and 7 T (Figure 5B), we found that the measured 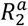 values at the two field strengths are comparable within the limits of precision: 6.88 ± 0.32 s^-1^ at 3 T versus 7.05 ± 1.26 s^-1^ at 7 T. Furthermore, no significant differences were observed between 3 T (blue) and 7 T (red) in a separate study from the literature [42], where 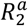 values were derived from specific fiber tracts. For example, in the corpus callosum (CC, Figure 5C), the values were 10.98 ± 0.74 s^-1^ at 3 T and 9.42 ± 2.91 s^-1^ at 7 T; in the cingulum (CG, Figure 5D), they were 11.03 ± 1.96 s^-1^ at 3 T and 10.22 ± 2.05 s^-1^ at 7 T. These findings suggest that susceptibility effects are not the dominant mechanism underlying anisotropic transverse relaxation, contrary to previous assumptions [3, 7, 8], and consistent with the conclusions discussed above.

**FIGURE 5.**
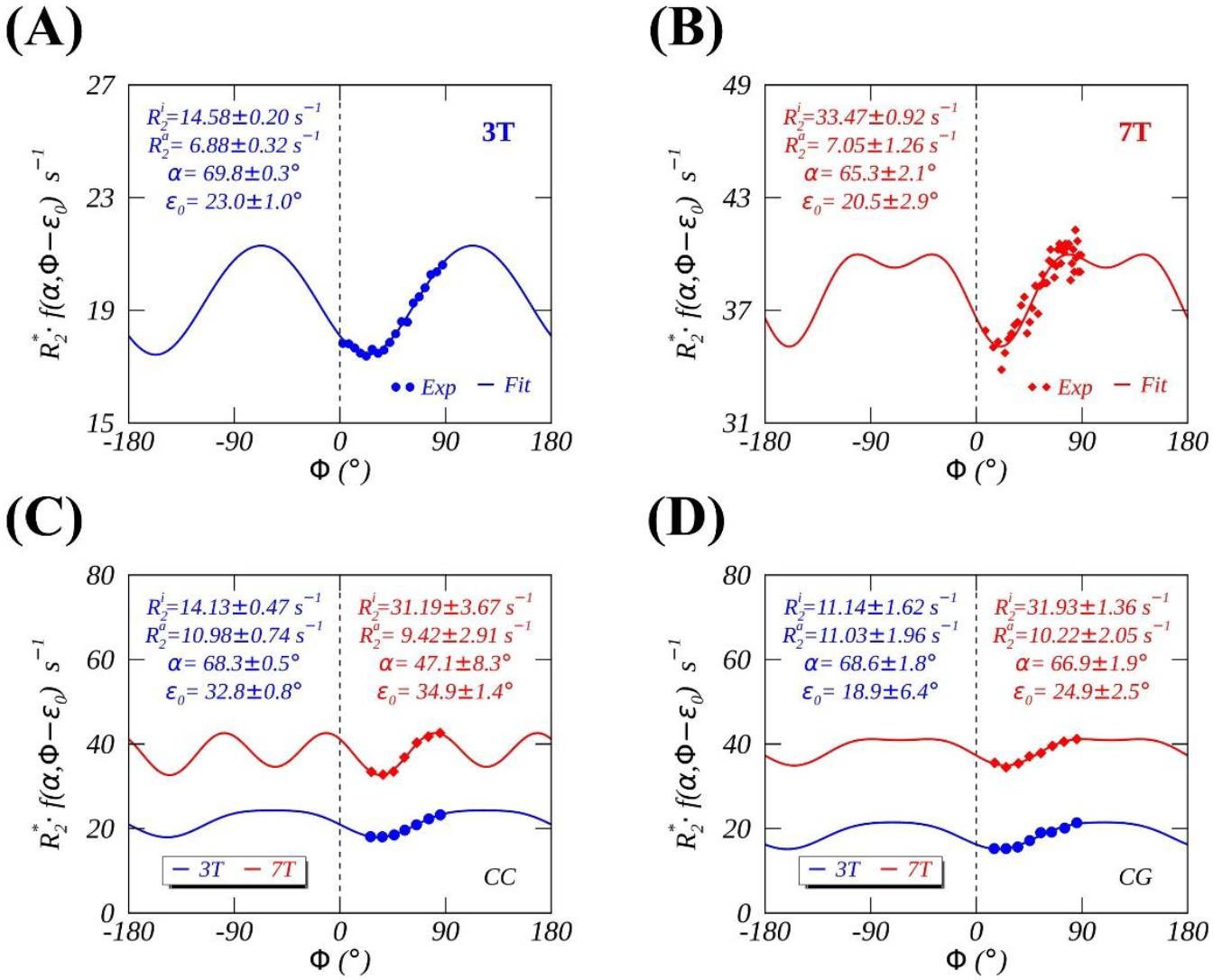
Measured (colored symbols) and fitted (colored lines) orientation-dependent transverse relaxation rates 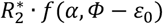 at 3 T (blue lines) and 7 T (red lines). Orientation-dependent 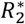 profiles were derived from: (A, B) whole brain WM; (C) corpus callosum; and (D) cingulum.

## 5 DISCUSSION

The aim of this study is to elucidate the long-standing challenges surrounding the origins of orientation-dependent transverse relaxation in human brain WM. By comparing anisotropic transverse relaxation rates observed at 3 T and 7 T using both SE and GRE signals, we disentangled the respective contributions of MAE and susceptibility effects by leveraging their distinct dependences on the main magnetic field strength.

### 5.1 Distinct 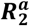 values obtained from SE and GRE measurements

Previously reported 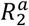 values in human brain WM have exhibited considerable variability across studies. This variability arises not only from differences in data acquisition protocols, such as pulse sequence design, echo spacing, and number of echoes, but also from variations in post-processing strategies, including signal fitting models and region-of-interest selection. For instance [24], Gil et al. demonstrated that 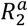 values derived from multi-echo SE (MESE) signals at 3 T were significantly lower than those obtained from multi-echo GRE (MEGE) signals. Specifically, MESE-based 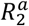 values ranged from 0 to 1.5 s^-1^, while MEGE-based values ranged from 3.1 to 4.5 s^-1^, underscoring the influence of sequence type on the estimation of anisotropic transverse relaxation.

In the present study, we observed a similar trend. The average 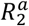 value derived from single SE signals embedded in DTI datasets was approximately 4.0 s^-1^ (Table S1), which is notably lower than the GRE-derived values of around 7.0 s^-1^ (Figures 5A-5B) for whole-brain WM at both 3 T and 7 T. Moreover, GRE-based values were even higher when the analysis was focused on specific fiber bundles, such as the corpus callosum (Figure 5C) and the cingulum (Figure 5D), indicating that these specific fiber tracts exhibit relatively higher microstructural anisotropies and orientation-dependent relaxation effects.

To mitigate methodological inconsistencies and isolate the influence of magnetic field strength, we restricted our comparison of 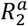 values to datasets acquired using similar pulse sequences and post-processing workflows at both 3 T and 7 T. This approach minimized confounding factors and ensured that the observed differences in 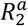 values primarily reflect true effects associated with field strengths, rather than artifacts introduced by acquisition or analysis variability.

While *B*_1_ field inhomogeneity becomes inevitable across the whole brain at higher field strengths, our analysis focuses exclusively on a subset of WM voxels exhibiting highly organized (0.5 < FA < 0.9 and 0.5 < MO < 1.0) microstructure. These selected voxels, derived from various brain regions, were binned and averaged according to their orientations to generate orientation-dependent *R*_2_ profiles. This customized processing workflow is expected to reduce the potential impact of *B*_1_ inhomogeneity on our results.

### 5.2 *B*_0_-dependent 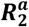 from SE signals

Based on single SE signals acquired from whole-brain WM in 25 subjects, we observed an average increase in 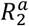 from 3 T to 7 T of 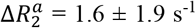, indicating a modest, though not dominant, as previously assumed, contribution of susceptibility effects to the measured 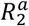at 3 T. This contribution was estimated at 8.3 ± 10.2%, suggesting that while susceptibility effects are present, they account for only a minor portion of the anisotropic transverse relaxation at this field strength.

This finding is consistent with prior results from *B*_0_-dependent *T*_1ρ_ dispersion studies in rat brain WM [5, 48], in which susceptibility effects accounted for approximately 12% of the transverse relaxation dispersion at 3 T [5], with the remaining contrast attributed to MAE. Supporting this interpretation, Bauser et al. recently reported negligible changes in orientation-dependent *R*_2_ of single-fiber WM voxels between 1.5 T and 3 T using MESE measurements [47], further reinforcing the notion that MAE is the dominant mechanism driving anisotropic transverse relaxation at lower field strengths.

It is worth emphasizing, however, that Bauser et al. did not identify the dominant relaxation mechanism. Instead, they merely state that the field-strength dependence of relaxation anisotropy reflects multiple contributing sources in WM fibers [47]. This inconclusive statement may lead prospective readers to incorrectly infer that a susceptibility-based mechanism governs the observed orientation-dependent *R*_2_ behavior.

Average *T*_2_ (i.e., 1/*R*_2_) values in WM have also been reported across field strengths ranging from 0.35 T to 9.4 T in two healthy volunteers [49]. When comparing *T*_2_ values at 3 T (62 ± 2 ms) and 7 T (37 ± 3 ms), the susceptibility-related contribution at 3 T is approximately 14%, assuming that the decrease in *T*_2_ with increasing field strength is attributed exclusively to *orientation-dependent* susceptibility-based relaxation pathway. A similar comparison between 3 T and 9.4 T (29 ± 2 ms) yields a susceptibility-related contribution at 3 T of approximately 12%.

To our knowledge, this study represents the first in vivo characterization of orientation-dependent *R*_2_ profiles in human brain WM at 7 T using single SE signals. Previous investigations using formalin-fixed brain samples at 7 T reported minimal orientation dependence in MESE-based *R*_2_ compared to GRE-based 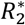, leading to the conclusion that susceptibility effects were the primary source of anisotropic transverse relaxation [3]. However, temperature-dependent 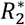 anisotropies observed in the same study pointed to MAE as a more plausible contributor, suggesting that the challenges [50] associated with accurately measuring *R*_2_, particularly at 7 T [51], may obscure the true biophysical origins of orientation-dependent transverse relaxation in WM.

### 5.3 *B*_0_-dependent 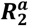 from GRE signals

The conclusion regarding the dominant mechanism of anisotropic transverse relaxation in brain WM at 3 T is further supported by GRE-based 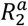 values reported in both the present study and previous investigations across different *B*_0_ field strengths. Notably, this conclusion is strengthened by the consistency of GRE-based 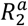 values observed across studies conducted at varying field strengths. Despite differences in data acquisition and analysis methodologies between two independent studies conducted by separate research groups [40, 41], the GRE-based 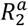 values derived from whole-brain WM at both 3 T (Figure 5A) and 7 T (Figure 5B) in the present study were consistently around 7 s^-1^. In both investigations, the adverse FAE on orientation-dependent 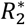 profiles were mitigated through linear regression across multiple profiles acquired with different flip angles, as detailed in the literature [41].

When the analysis was focused on specific fiber tracts, such as the corpus callosum (Figure 5C) and the cingulum (Figure 5D) [42], the 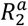 values increased to approximately 10 s^-1^. However, these elevated values remained statistically indistinguishable between the two *B*_0_ field strengths, suggesting that the observed increases are primarily attributable to relatively higher microstructural anisotropies in these regions. These findings further support the notion that MAE, rather than susceptibility effects, is the dominant contributor to anisotropic transverse relaxation in WM at 3 T.

Comparable GRE-based orientation-dependent *R*_2_ values have been reported across a wide range of *B*_0_ field strengths in a substantial body of literature [13, 24, 25]. In many of these studies, the metric 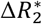, defined as the difference in 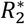 between perpendicular and parallel axonal orientations, is used as a proxy for 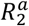. These values typically range between 3 and 5 s^-1^. For instance, Rudko et al. [52] reported a 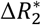 of approximately 4 s^-1^ in fixed rat brain at 9.4 T, while Wharton et al. [7] found similar values in fresh pig brain at 7 T. In both cases, orientation information was obtained by physically rotating brain samples within the magnet, rather than relying on principal diffusivity directions derived from DTI, thereby providing a more direct assessment of 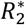 orientation dependence.

It is important to note that direct comparisons between the GRE-based 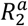 values reported in the present study and previously published 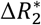 values are not strictly meaningful, as the same orientation-dependent 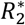 profile can yield different numerical values depending on the metric used. Nonetheless, the consistent observation of comparable 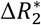 values across different species, tissue preparations, and magnetic field strengths provides compelling evidence that MAE is the dominant mechanism underlying anisotropic transverse relaxation at 3 T in brain WM. This conclusion is further supported by recent advances in orientation-dependent relaxation theory [25], which have effectively ruled out susceptibility effects as a plausible primary mechanism for anisotropic transverse relaxation in WM.

## 6 CONCLUSIONS

This study identifies MAE as the predominant mechanism underlying orientation-dependent transverse relaxation in human brain WM at 3 T, offering a distinct perspective from earlier interpretations in the literature. By disentangling the respective contributions of susceptibility effects and MAE across magnetic field strengths, our findings provide compelling evidence that MAE, rather than susceptibility effects, is the primary drive of the observed anisotropic transverse relaxation behavior at lower fields. These insights carry important implications for advancing the biophysical understanding of anisotropic relaxation contrast in MRI and contribute to a more accurate interpretation of WM microstructure in both clinical and research settings.

## DATA AVAILABILITY STATEMENT

Data and Python/IDL scripts used in this work are available upon request from the authors.

## Supporting Information

**TABLE S1.**
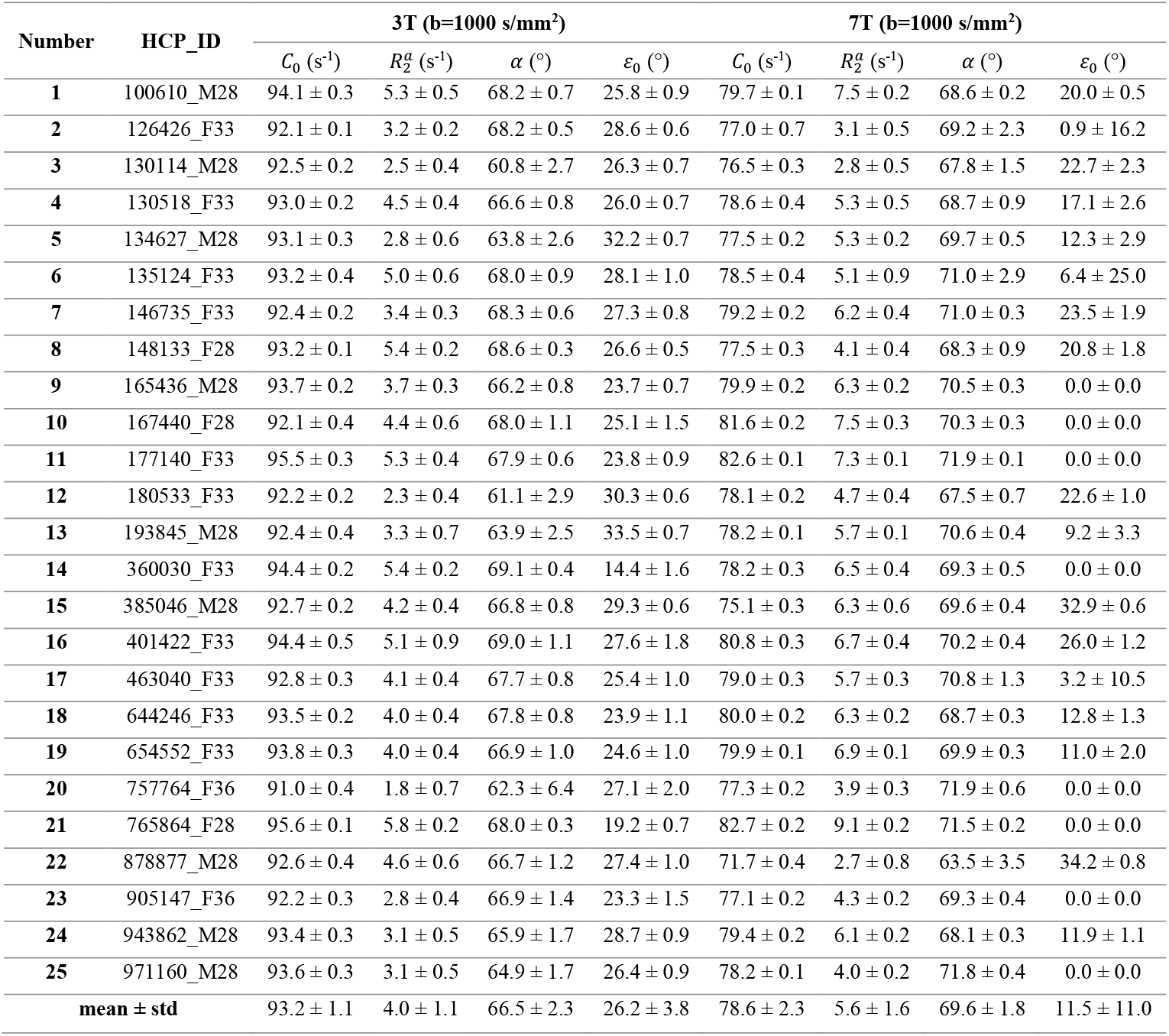
Fitted parameters from the orientation-dependent transverse relaxation model for 25 subjects, based on DTI datasets acquired with a non-zero b-value of 1000 s/mm^2^ at 3 T and 7 T. Values are presented as mean ± standard deviation.

| Number | HCP_ID | 3T (b=1000 s/mm<sup>2</sup>) |  |  |  | 7T (b=1000 s/mm<sup>2</sup>) |  |  |  |
| --- | --- | --- | --- | --- | --- | --- | --- | --- | --- |
| | | $C_0$ (s<sup>-1</sup>) | $R_2^a$ (s<sup>-1</sup>) | $\alpha$ (°) | $\varepsilon_0$ (°) | $C_0$ (s<sup>-1</sup>) | $R_2^a$ (s<sup>-1</sup>) | $\alpha$ (°) | $\varepsilon_0$ (°) |
| 1 | 100610_M28 | 94.1 ± 0.3 | 5.3 ± 0.5 | 68.2 ± 0.7 | 25.8 ± 0.9 | 79.7 ± 0.1 | 7.5 ± 0.2 | 68.6 ± 0.2 | 20.0 ± 0.5 |
| 2 | 126426_F33 | 92.1 ± 0.1 | 3.2 ± 0.2 | 68.2 ± 0.5 | 28.6 ± 0.6 | 77.0 ± 0.7 | 3.1 ± 0.5 | 69.2 ± 2.3 | 0.9 ± 16.2 |
| 3 | 130114_M28 | 92.5 ± 0.2 | 2.5 ± 0.4 | 60.8 ± 2.7 | 26.3 ± 0.7 | 76.5 ± 0.3 | 2.8 ± 0.5 | 67.8 ± 1.5 | 22.7 ± 2.3 |
| 4 | 130518_F33 | 93.0 ± 0.2 | 4.5 ± 0.4 | 66.6 ± 0.8 | 26.0 ± 0.7 | 78.6 ± 0.4 | 5.3 ± 0.5 | 68.7 ± 0.9 | 17.1 ± 2.6 |
| 5 | 134627_M28 | 93.1 ± 0.3 | 2.8 ± 0.6 | 63.8 ± 2.6 | 32.2 ± 0.7 | 77.5 ± 0.2 | 5.3 ± 0.2 | 69.7 ± 0.5 | 12.3 ± 2.9 |
| 6 | 135124_F33 | 93.2 ± 0.4 | 5.0 ± 0.6 | 68.0 ± 0.9 | 28.1 ± 1.0 | 78.5 ± 0.4 | 5.1 ± 0.9 | 71.0 ± 2.9 | 6.4 ± 25.0 |
| 7 | 146735_F33 | 92.4 ± 0.2 | 3.4 ± 0.3 | 68.3 ± 0.6 | 27.3 ± 0.8 | 79.2 ± 0.2 | 6.2 ± 0.4 | 71.0 ± 0.3 | 23.5 ± 1.9 |
| 8 | 148133_F28 | 93.2 ± 0.1 | 5.4 ± 0.2 | 68.6 ± 0.3 | 26.6 ± 0.5 | 77.5 ± 0.3 | 4.1 ± 0.4 | 68.3 ± 0.9 | 20.8 ± 1.8 |
| 9 | 165436_M28 | 93.7 ± 0.2 | 3.7 ± 0.3 | 66.2 ± 0.8 | 23.7 ± 0.7 | 79.9 ± 0.2 | 6.3 ± 0.2 | 70.5 ± 0.3 | 0.0 ± 0.0 |
| 10 | 167440_F28 | 92.1 ± 0.4 | 4.4 ± 0.6 | 68.0 ± 1.1 | 25.1 ± 1.5 | 81.6 ± 0.2 | 7.5 ± 0.3 | 70.3 ± 0.3 | 0.0 ± 0.0 |
| 11 | 177140_F33 | 95.5 ± 0.3 | 5.3 ± 0.4 | 67.9 ± 0.6 | 23.8 ± 0.9 | 82.6 ± 0.1 | 7.3 ± 0.1 | 71.9 ± 0.1 | 0.0 ± 0.0 |
| 12 | 180533_F33 | 92.2 ± 0.2 | 2.3 ± 0.4 | 61.1 ± 2.9 | 30.3 ± 0.6 | 78.1 ± 0.2 | 4.7 ± 0.4 | 67.5 ± 0.7 | 22.6 ± 1.0 |
| 13 | 193845_M28 | 92.4 ± 0.4 | 3.3 ± 0.7 | 63.9 ± 2.5 | 33.5 ± 0.7 | 78.2 ± 0.1 | 5.7 ± 0.1 | 70.6 ± 0.4 | 9.2 ± 3.3 |
| 14 | 360030_F33 | 94.4 ± 0.2 | 5.4 ± 0.2 | 69.1 ± 0.4 | 14.4 ± 1.6 | 78.2 ± 0.3 | 6.5 ± 0.4 | 69.3 ± 0.5 | 0.0 ± 0.0 |
| 15 | 385046_M28 | 92.7 ± 0.2 | 4.2 ± 0.4 | 66.8 ± 0.8 | 29.3 ± 0.6 | 75.1 ± 0.3 | 6.3 ± 0.6 | 69.6 ± 0.4 | 32.9 ± 0.6 |
| 16 | 401422_F33 | 94.4 ± 0.5 | 5.1 ± 0.9 | 69.0 ± 1.1 | 27.6 ± 1.8 | 80.8 ± 0.3 | 6.7 ± 0.4 | 70.2 ± 0.4 | 26.0 ± 1.2 |
| 17 | 463040_F33 | 92.8 ± 0.3 | 4.1 ± 0.4 | 67.7 ± 0.8 | 25.4 ± 1.0 | 79.0 ± 0.3 | 5.7 ± 0.3 | 70.8 ± 1.3 | 3.2 ± 10.5 |
| 18 | 644246_F33 | 93.5 ± 0.2 | 4.0 ± 0.4 | 67.8 ± 0.8 | 23.9 ± 1.1 | 80.0 ± 0.2 | 6.3 ± 0.2 | 68.7 ± 0.3 | 12.8 ± 1.3 |
| 19 | 654552_F33 | 93.8 ± 0.3 | 4.0 ± 0.4 | 66.9 ± 1.0 | 24.6 ± 1.0 | 79.9 ± 0.1 | 6.9 ± 0.1 | 69.9 ± 0.3 | 11.0 ± 2.0 |
| 20 | 757764_F36 | 91.0 ± 0.4 | 1.8 ± 0.7 | 62.3 ± 6.4 | 27.1 ± 2.0 | 77.3 ± 0.2 | 3.9 ± 0.3 | 71.9 ± 0.6 | 0.0 ± 0.0 |
| 21 | 765864_F28 | 95.6 ± 0.1 | 5.8 ± 0.2 | 68.0 ± 0.3 | 19.2 ± 0.7 | 82.7 ± 0.2 | 9.1 ± 0.2 | 71.5 ± 0.2 | 0.0 ± 0.0 |
| 22 | 878877_M28 | 92.6 ± 0.4 | 4.6 ± 0.6 | 66.7 ± 1.2 | 27.4 ± 1.0 | 71.7 ± 0.4 | 2.7 ± 0.8 | 63.5 ± 3.5 | 34.2 ± 0.8 |
| 23 | 905147_F36 | 92.2 ± 0.3 | 2.8 ± 0.4 | 66.9 ± 1.4 | 23.3 ± 1.5 | 77.1 ± 0.2 | 4.3 ± 0.2 | 69.3 ± 0.4 | 0.0 ± 0.0 |
| 24 | 943862_M28 | 93.4 ± 0.3 | 3.1 ± 0.5 | 65.9 ± 1.7 | 28.7 ± 0.9 | 79.4 ± 0.2 | 6.1 ± 0.2 | 68.1 ± 0.3 | 11.9 ± 1.1 |
| 25 | 971160_M28 | 93.6 ± 0.3 | 3.1 ± 0.5 | 64.9 ± 1.7 | 26.4 ± 0.9 | 78.2 ± 0.1 | 4.0 ± 0.2 | 71.8 ± 0.4 | 0.0 ± 0.0 |
| mean ± std | | 93.2 ± 1.1 | 4.0 ± 1.1 | 66.5 ± 2.3 | 26.2 ± 3.8 | 78.6 ± 2.3 | 5.6 ± 1.6 | 69.6 ± 1.8 | 11.5 ± 11.0 |

**TABLE S2.** Fitted parameters from the orientation-dependent transverse relaxation model for 25 subjects, based on DTI datasets acquired with a non-zero b-value of 2000 s/mm^2^ at 3 T and 7 T. Values are reported as mean ± standard deviation.

| Number | HCP_ID | 3T (b=2000 s/mm<sup>2</sup>) |  |  |  | 7T (b=2000 s/mm<sup>2</sup>) |  |  |  |
| --- | --- | --- | --- | --- | --- | --- | --- | --- | --- |
| | | $C_0$ (s<sup>-1</sup>) | $R_2^a$ (s<sup>-1</sup>) | $\alpha$ (°) | $\epsilon_0$ (°) | $C_0$ (s<sup>-1</sup>) | $R_2^a$ (s<sup>-1</sup>) | $\alpha$ (°) | $\epsilon_0$ (°) |
| 1 | 100610_M28 | 94.8 ± 0.4 | 6.2 ± 0.7 | 69.1 ± 0.7 | 27.1 ± 1.2 | 80.6 ± 0.1 | 8.3 ± 0.2 | 69.6 ± 0.2 | 21.5 ± 0.6 |
| 2 | 126426_F33 | 92.3 ± 0.3 | 3.4 ± 0.4 | 68.9 ± 0.8 | 29.8 ± 1.1 | 76.9 ± 0.2 | 2.6 ± 0.3 | 69.1 ± 1.0 | 0.0 ± 0.0 |
| 3 | 130114_M28 | 92.8 ± 0.2 | 3.0 ± 0.4 | 62.1 ± 1.8 | 28.5 ± 0.5 | 76.8 ± 0.4 | 2.5 ± 0.7 | 66.5 ± 2.2 | 33.8 ± 1.0 |
| 4 | 130518_F33 | 93.4 ± 0.3 | 5.1 ± 0.6 | 67.6 ± 0.9 | 26.9 ± 0.9 | 78.4 ± 0.3 | 4.8 ± 0.4 | 68.9 ± 0.8 | 18.7 ± 2.4 |
| 5 | 134627_M28 | 93.3 ± 0.4 | 2.9 ± 0.8 | 63.5 ± 3.3 | 34.2 ± 0.8 | 78.0 ± 0.1 | 5.2 ± 0.5 | 71.4 ± 1.0 | 7.1 ± 8.5 |
| 6 | 135124_F33 | 92.9 ± 0.4 | 4.3 ± 0.7 | 67.1 ± 1.3 | 29.3 ± 1.0 | 78.2 ± 0.5 | 4.4 ± 0.9 | 70.9 ± 3.6 | 5.4 ± 30.9 |
| 7 | 146735_F33 | 92.4 ± 0.2 | 3.2 ± 0.4 | 67.8 ± 0.9 | 30.3 ± 0.8 | 79.4 ± 0.2 | 6.1 ± 0.5 | 71.7 ± 0.4 | 19.4 ± 4.0 |
| 8 | 148133_F28 | 93.5 ± 0.2 | 5.8 ± 0.3 | 69.2 ± 0.3 | 26.6 ± 0.5 | 77.6 ± 0.2 | 4.0 ± 0.3 | 69.5 ± 0.7 | 19.8 ± 2.4 |
| 9 | 165436_M28 | 93.9 ± 0.2 | 3.8 ± 0.4 | 66.6 ± 1.0 | 25.1 ± 0.9 | 79.7 ± 0.2 | 5.7 ± 0.2 | 70.7 ± 0.3 | 0.0 ± 0.0 |
| 10 | 167440_F28 | 92.4 ± 0.5 | 4.8 ± 0.8 | 68.5 ± 1.2 | 26.1 ± 1.8 | 81.5 ± 0.2 | 7.3 ± 0.3 | 70.5 ± 0.3 | 0.0 ± 0.0 |
| 11 | 177140_F33 | 95.4 ± 0.3 | 5.0 ± 0.4 | 67.6 ± 0.7 | 25.1 ± 0.8 | 82.1 ± 0.1 | 6.3 ± 0.2 | 71.7 ± 0.2 | 0.0 ± 0.0 |
| 12 | 180533_F33 | 92.5 ± 0.2 | 2.5 ± 0.4 | 62.5 ± 2.5 | 31.5 ± 0.6 | 78.9 ± 0.3 | 5.6 ± 0.4 | 69.1 ± 0.6 | 22.3 ± 1.5 |
| 13 | 193845_M28 | 92.4 ± 0.4 | 3.0 ± 0.7 | 61.7 ± 3.7 | 35.0 ± 0.6 | 78.4 ± 0.1 | 5.4 ± 0.3 | 71.9 ± 0.4 | 5.9 ± 3.5 |
| 14 | 360030_F33 | 94.6 ± 0.1 | 5.4 ± 0.2 | 69.5 ± 0.3 | 14.5 ± 1.6 | 77.8 ± 0.3 | 5.0 ± 0.5 | 69.3 ± 0.8 | 0.0 ± 0.0 |
| 15 | 385046_M28 | 93.6 ± 0.4 | 5.0 ± 0.6 | 67.7 ± 0.9 | 30.6 ± 0.7 | 76.5 ± 0.4 | 8.0 ± 0.8 | 70.6 ± 0.3 | 32.6 ± 0.9 |
| 16 | 401422_F33 | 94.6 ± 0.6 | 5.2 ± 1.0 | 69.2 ± 1.1 | 28.5 ± 1.9 | 81.1 ± 0.3 | 6.5 ± 0.5 | 71.1 ± 0.3 | 25.1 ± 1.9 |
| 17 | 463040_F33 | 92.8 ± 0.3 | 3.8 ± 0.5 | 66.9 ± 1.1 | 27.5 ± 0.9 | 78.8 ± 0.3 | 5.2 ± 0.4 | 70.9 ± 1.6 | 2.0 ± 13.2 |
| 18 | 644246_F33 | 93.6 ± 0.3 | 3.9 ± 0.5 | 67.9 ± 1.0 | 25.3 ± 1.3 | 80.0 ± 0.2 | 5.9 ± 0.2 | 69.4 ± 0.5 | 9.8 ± 2.6 |
| 19 | 654552_F33 | 94.3 ± 0.3 | 4.6 ± 0.5 | 67.7 ± 0.9 | 27.2 ± 1.0 | 80.3 ± 0.1 | 6.5 ± 0.3 | 71.5 ± 0.5 | 2.7 ± 3.9 |
| 20 | 757764_F36 | 91.0 ± 0.5 | 1.2 ± 0.4 | 49.6 ± 14.0 | 29.6 ± 2.1 | 77.3 ± 0.2 | 3.6 ± 0.3 | 71.8 ± 0.8 | 0.0 ± 0.0 |
| 21 | 765864_F28 | 95.9 ± 0.1 | 6.1 ± 0.2 | 68.3 ± 0.3 | 20.3 ± 0.6 | 82.4 ± 0.1 | 8.4 ± 0.2 | 72.0 ± 0.2 | 0.0 ± 0.0 |
| 22 | 878877_M28 | 93.0 ± 0.3 | 4.6 ± 0.5 | 66.2 ± 0.9 | 30.4 ± 0.5 | 72.7 ± 0.7 | 3.7 ± 1.2 | 67.0 ± 2.4 | 35.2 ± 1.2 |
| 23 | 905147_F36 | 92.5 ± 0.3 | 3.1 ± 0.5 | 68.0 ± 1.2 | 24.8 ± 1.6 | 77.2 ± 0.2 | 4.1 ± 0.3 | 68.9 ± 0.6 | 0.0 ± 0.0 |
| 24 | 943862_M28 | 93.9 ± 0.4 | 3.9 ± 0.7 | 65.9 ± 1.7 | 32.6 ± 0.7 | 80.0 ± 0.3 | 6.2 ± 0.3 | 69.5 ± 0.7 | 13.0 ± 3.3 |
| 25 | 971160_M28 | 93.8 ± 0.3 | 3.2 ± 0.5 | 65.3 ± 1.5 | 27.8 ± 0.8 | 78.1 ± 0.2 | 3.7 ± 0.5 | 72.1 ± 1.0 | 5.4 ± 7.6 |
| mean ± std | | 93.4 ± 1.1 | 4.1 ± 1.2 | 66.2 ± 4.0 | 27.8 ± 4.1 | 78.7 ± 2.1 | 5.4 ± 1.6 | 70.2 ± 1.5 | 11.2 ± 11.7 |

